# Direct Observation of Electrically Conductive Pili Emanating from *Geobacter sulfurreducens*

**DOI:** 10.1101/2021.07.06.451359

**Authors:** Xinying Liu, David J. F. Walker, Stephen S. Nonnenmann, Dezhi Sun, Derek R. Lovley

**Affiliations:** Department of Microbiology, University of Massachusetts—Amherst, Amherst, Massachusetts, USA; College of Environmental Science and Engineering, Beijing Forestry University, Beijing, 100083, China; Institute for Cellular and Molecular Biology, University of Texas at Austin, Austin, Texas 78712, USA; Institute for Applied Life Sciences, university of Massachusetts—Amherst, Amherst, Massachusetts, USA; Department of Mechanical and Industrial Engineering, University of Massachusetts—Amherst, Amherst, Massachusetts, USA

## Abstract

*Geobacter sulfurreducens* is a model microbe for elucidating the mechanisms for extracellular electron transfer in several biogeochemical cycles, bioelectrochemical applications, and microbial metal corrosion. Multiple lines of evidence previously suggested that electrically conductive pili (e-pili) are an essential conduit for long-range extracellular electron transport in *G. sulfurreducens*. However, it has recently been reported that *G. sulfurreducens* does not express e-pili and that filaments comprised of multi-heme *c*-type cytochromes are responsible for long-range electron transport. This possibility was directly investigated by examining cells, rather than filament preparations, with atomic force microscopy. Approximately 90 % of the filaments emanating from wild-type cells had a diameter (3 nm) and conductance consistent with previous reports of e-pili harvested from *G. sulfurreducens* or heterologously expressed in *E. coli* from the *G. sulfurreducens* pilin gene. The remaining 10% of filaments had a morphology consistent with filaments comprised of the *c*-type cytochrome OmcS. A strain expressing a modified pilin gene designed to yield poorly conductive pili expressed 90 % filaments with a 3 nm diameter, but greatly reduced conductance, further indicating that the 3 nm diameter conductive filaments in the wild-type strain were e-pili. A strain in which genes for five of the most abundant outer-surface *c*-type cytochromes, including OmcS, was deleted yielded only 3 nm diameter filaments with the same conductance as in the wild-type. These results demonstrate that e-pili are the most abundant conductive filaments expressed by *G. sulfurreducens*, consistent with previous functional studies demonstrating the need for e-pili for long-range extracellular electron transfer.

**Importance:** Electroactive microbes have significant environmental impacts as well as applications in bioenergy and bioremediation. The composition, function, and even existence of electrically conductive pili (e-pili) has been one of the most contentious areas of investigation in electromicrobiology, in part because e-pili offer a mechanism for long-range electron transport that does not involve the metal co-factors common in much of biological electron transport. This study demonstrates that e-pili are abundant filaments emanating from *Geobacter sulfurreducens*, which serves as a model for long-range extracellular electron transfer in direct interspecies electron transfer, dissimilatory metal reduction, microbe-electrode exchange, and corrosion caused by direct electron uptake from Fe(0). The methods described in this study provide a simple strategy for evaluating the distribution of conductive filaments throughout the microbial world with an approach that avoids artifactual production and/or enrichment of filaments that may not be physiologically relevant.

---

Electroactive microorganisms are important in multiple biogeochemical cycles, the human gut, several bioenergy strategies, and metal corrosion (1, 2). One of the most contentious issues in electromicrobiology has been the role of electrically conductive protein nanowires in facilitating long-range electron transport. Electrically conductive protein nanowires have been studied most extensively in *Geobacter sulfurreducen*s, which has served as the model microbe for elucidating the mechanisms of long-range electron transport in *Geobacter* species (3). *Geobacter* are of interest because they are often the most abundant electroactive microbes in soils and sediments in which organic matter oxidation is coupled to Fe(III) oxide reduction; in natural methanogenic environments and anaerobic digesters where they serve as electron-donating partners for direct interspecies electron transfer (DIET) with methanogens; and in electrode biofilms harvesting electricity from waste organic matter (3–5). Furthermore, *Geobacter* are the most effective microbes available in culture for extracellular electron transport functions such as Fe(III) oxide reduction (3), producing electric current (5), DIET (6), and corrosion via direct extraction of electrons from metallic iron (7, 8). An additional area of interest is the potential for constructing electronic devices with novel functions with *G. sulfurreducens* protein nanowires (9).

Debate has arisen over the composition of *G. sulfurreducens* protein nanowires and their role in long-range electron transfer. Multiple lines of evidence have suggested that electrically conductive pili (e-pili) are the most abundant *G. sulfurreducens* protein nanowires and that e-pili are essential for long-range electron transport (10, 11). However, two recent publications have suggested that *G. sulfurreducens* does not express e-pili and that protein nanowires comprised of the multi-heme *c-*type cytochromes OmcS and OmcZ are the functional conduits for long-range extracellular electron transfer (12, 13). The primary argument against the production of e-pili is the fact filaments comprised of *c*-type cytochromes are the most abundant filaments observed in filament preparations observed with cryo-electron microscopy (12, 13). However, generating these filament preparations involves shearing filaments from the cell, purifying the filaments under high pH, selective precipitation with ammonium sulfate, and affixing filaments to grids. Each of these steps has the potential to selectively enrich specific filaments or for artifactual formation of cytochrome filaments (11). For example, in studies of *G. sulfurreducens* filament preparations prepared by the same person in the same laboratory, under identical conditions, cryo-electron microscopy suggest that a majority of the filaments were comprised of OmcS (14), whereas filaments with a diameter consistent with e-pili, not OmcS, were observed with atomic force microscopy (AFM) (15).

## Direct AFM of Cells

In order to avoid potential artifacts/enrichments associated with filament purification, the filaments associated with *G. sulfurreducens* cells were directly examined with AFM. To simplify the analysis, cells were grown with fumarate as the electron acceptor, a growth condition in which the pilin monomer PilA and OmcS are expressed, but expression of the gene for the multi-heme cytochrome OmcZ is repressed (16–18). AFM of culture aliquots directly deposited on a conductive surface revealed cells with abundant filaments (Fig. 1A and Supplemental Fig. S1A). There were two types of filaments emanating from the cells. One filament type appeared to be comprised of OmcS, as evidenced from its 4 nm diameter (Fig. 1B and Supplemental Fig. S1B) and its characteristic axial periodicity with a 20 nm pitch (12, 14) (Fig. 1C). The OmcS filaments consistently accounted for only ca. 10 % of the filaments observed (Fig.1A and Supplemental Figs. S2-4).

**Fig. 1.**
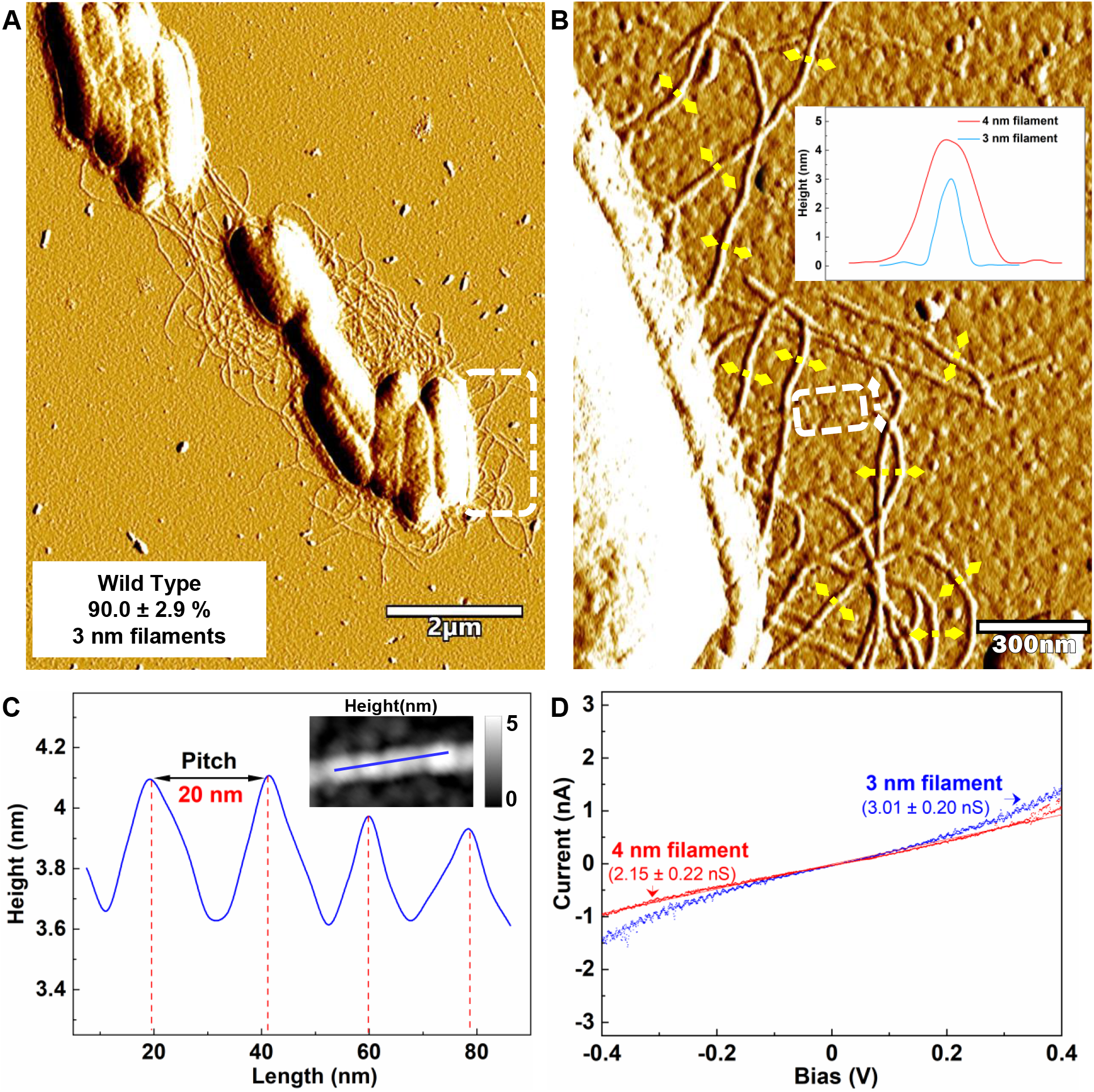
Characterization of filaments emanating from G. sulfurreducens with the wild-type pilin gene. (A) AFM amplitude image. The proportion of 3 nm diameter filaments was calculated from the total number of 3 nm and 4 nm diameter filaments counted in 9 regions from 3 separate samples (Supplemental Figs. 2-4) and were determined from height images similar to those shown in Supplemental Fig. 1. (B) Higher magnification of the region highlighted in the dashed frame in panel A. Inset shows typical height profiles across the 3 nm (yellow lines) and 4 nm (white line) diameter filaments, as determined from the corresponding height images (Supplemental Fig. S1B). Due to fluctuation of diameter along the axis of the filaments, diameters were measured at the points of greatest diameter for consistency. (C) Longitudinal height profile (along solid blue line in inset) for region on the 4 nm filament noted by the white dashed frame in panel B. (D) Comparison of point-mode current response (I-V) spectroscopy for 4 nm (red) and 3 nm (blue) diameter filaments. The responses shown are representative of three different measurements on each of three individual filaments (Supplemental Figs. S5 and S6). Conductance (mean + standard deviation, n=9) was calculated from a linear fit model between −0.2 V and 0.2 V (Supplemental Figs. S5 and S6).

Approximately 90% of the filaments were 3 nm in diameter (Fig. 1A and Supplemental Figs. S2-4), the same diameter as the filaments observed when the *G. sulfurreducens* PilA gene is expressed in *Pseudomonas aeruginosa* (19) or *Escherichia coli* (20) and the same diameter of individual conductive filaments previously harvested from *G. sulfurreducens* (16, 21). These results suggest that the 3 nm diameter filaments are e-pili. As expected from the growth conditions employed, no filaments with a morphology consistent with the 2.5 nm diameter and axial pitch of OmcZ filaments (13) were observed. Both the OmcS and e-pili filaments exhibited an ohmic-like response (Fig. 1D). The conductance of the e-pili was slightly higher than that of the OmcS filaments (Fig. 1D).

*G. sulfurreducens* strain Aro-5 was previously constructed to replace the PilA pilin gene with *aro-5*, a synthetic pilin gene designed to yield poorly conductive pili (18). The conductivity of filaments harvested from the cells is much lower than the conductivity of filaments harvested from wild-type controls (18, 21–23). Direct examination of filaments emanating from strain Aro-5 revealed two types of filaments, morphologically similar to those observed in the wild-type control (Figs. 2A, B and Supplemental Fig. S1C, D). Filaments with a diameter and longitudinal pitch (Fig. 2C) consistent with OmcS filaments comprised ca. 10 % of the filaments (Fig. 2A, Supplemental Figs. S7 and S8), similar to the OmcS filament abundance in the wild-type control and consistent with the observation that strain Aro-5 produces abundant OmcS (18). The conductance of these 4 nm diameter filaments was the same as the conductance observed for the OmcS filaments of the wild-type control (Fig. 2D and Supplemental Fig. S9). As with the wild-type strain, the 3 nm diameter filaments accounted for ca. 90% of the filaments observed, but their conductance was more than 100-fold lower (Fig. 2D and Supplemental Fig. S10). This decreased conductance is in agreement with previous observations of attenuated conductivity in filaments harvested from strain Aro-5, including measurements on individual 3 nm diameter filaments (18, 21–23). The dramatic change in the conductance of the 3 nm filaments emanating from cells associated with the expression of *aro-5* pilin gene provides further evidence that the 3 nm filaments in the wild-type strain were e-pili.

**Fig. 2.**
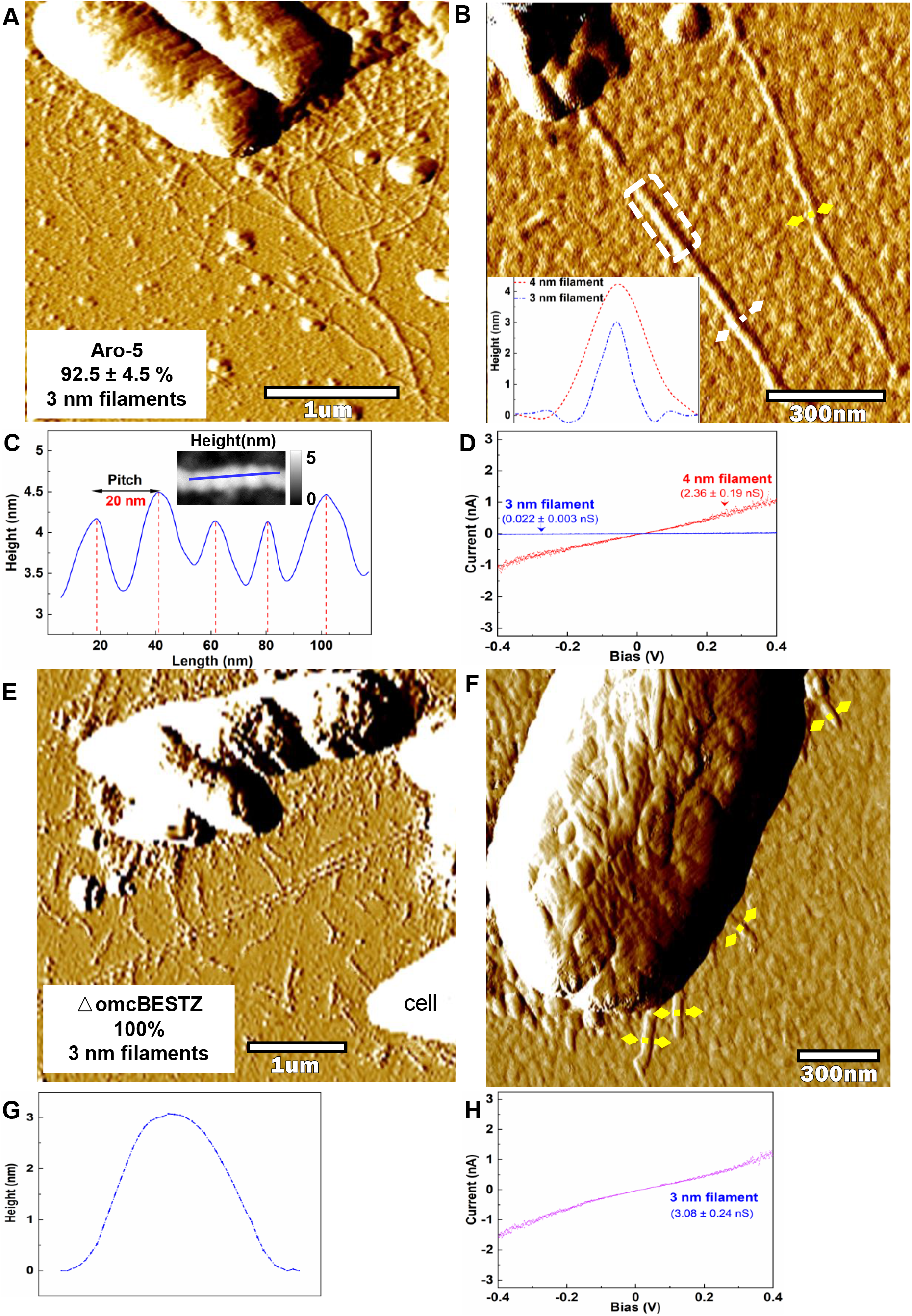
Characterization of filaments emanating from G. sulfurreducens strain Aro-5 and strain △omcBESTZ. (A) AFM amplitude image of filaments associated with strain Aro-5. The proportion of 3 nm diameter filaments was calculated from the total number of 3 nm and 4 nm diameter filaments counted in 6 regions from 3 separate samples (Supplemental Figs. S7 and S8) and were determined from height images similar to those shown in Supplemental Fig. 1. (B) AFM amplitude image at higher magnification illustrating the two filament types. Inset shows typical height profiles across the 3 nm (yellow lines) and 4 nm (white line) diameter filaments, as determined from the corresponding height images (Supplementary Fig. 1D). (C) Longitudinal height profile (along solid blue line in inset) for the portion of the 4 nm diameter filament within the white frame in panel B. (D) Comparison of point-mode current response (I-V) spectroscopy for 4 nm (red) and 3 nm (blue) filaments. The responses shown are representative of three different measurements on three individual wires (Supplemental Figs. S9 and S10). Conductance (mean + standard deviation, n=9) was calculated from a linear fit model between −0.2 V and 0.2 V (Supplemental Figs. S9 and S10). (E) AFM amplitude image of filaments associated with strain △omcBESTZ. (F) AFM amplitude image at higher magnification showing 3 nm diameter filaments emanating from cell of strain △omcBESTZ. (G) Typical height profile across the filaments designated by yellow lines in panel F, as determined from the corresponding height images (Supplementary Fig. 1F). (H) Point-mode current response (I-V) spectroscopy representative of three different measurements on three individual wires (Supplemental Fig. S11) on 3 nm filaments emanating from strain △omcBESTZ. Conductance (mean + standard deviation, n=9) was calculated from a linear fit model between −0.2 V and 0.2 V (Supplemental Fig. S11)..

In order to further investigate the possibility of cytochrome-based filaments, we next examined the previously described strain ΔomcBESTZ (24) in which the genes for the most abundant *G. sulfurreducens* outer surface multi-heme c-type cytochromes, OmcB, OmcE, OmcS, OmcT, and OmcZ were deleted. As expected, filaments with morphologies consistent with OmcS-based filaments were not apparent in this strain. All of the filaments emanating from strain ΔomcBESTZ and lying near the cells were short, but had a diameter of 3 nm (Fig. 2E, F and Supplemental Fig. S1E, F). Their conductance was the same as for the 3 nm filaments of the wild-type strain (Fig. 2H, Supplemental Fig. S11).

## Implications

The results of direct observation of filaments emanating from cells of *G. sulfurreducens* demonstrates that *G. sulfurreducens* copiously expresses filaments with properties expected for e-pili. The e-pili were ca. 10-fold more abundant than putative OmcS filaments. These observations are in accordance with a number of previous observations. For example, when a pilin monomer modified with a peptide tag was expressed in *G. sulfurreducens* all of the filaments observed emanating from the cells were also decorated with the peptide tag (25). Several studies reported recovery of electrically conductive 3 nm diameter filaments when filaments were sheared off the outer surface of *G. sulfurreducens* (16, 21, 25) or when the *G. sulfurreducens* pilin monomer was expressed in *P. aeruginosa* (19) or *E. coli* (20). Furthermore, as shown here, expressing *aro-5* instead of PilA resulted in 3 nm filaments emanating from the cells with a similar morphology, but greatly attenuated conductance. Heterologously expressing a pilin gene encoding increased aromatic amino acid content yielded 3 nm diameter filaments with 5000-fold higher conductivity than the wild-type control (26). These results are consistent with the expression of e-pili and inconsistent with cytochrome-based filaments, as was the finding reported here that the 3 nm filaments were still produced in a strain in which the genes for all the most abundant outer-surface cytochromes were deleted. The abundance of e-pili in *G. sulfurreducens* is also consistent with the finding that microbes that do not express outer-surface *c*-type cytochromes can construct conductive filaments from monomers homologous to the *G. sulfurreducens* pilin monomer (22, 23, 27).

Notably, *G. sulfurreducens* strains that express pili of low conductance are consistently deficient in long-range extracellular electron transfer (18, 22, 28), providing strong evidence for the role of e-pili in extracellular electron transport. The same cannot be said of the cytochrome filaments OmcS and OmcZ. *G. sulfurreducens* strain Aro-5 cannot produce highly conductive biofilms or high current densities on anodes (18), whereas deleting *omcS* has no impact on these phenotypes (17, 29). Deletion of *omcS* can inhibit Fe(III) oxide reduction, in some, but not all variants of *G. sulfurreducens* (30, 31). When deletion of *omcS* does have an impact, the strain can be rescued for Fe(III) oxide reduction with the addition of ultrafine-grained magnetite (32). However, magnetite cannot substitute for e-pili, demonstrating an essential role for e-pili in Fe(III) oxide reduction, but not for OmcS. OmcZ is not required for Fe(III) oxide reduction (17) and is not highly expressed in cells reducing Fe(III) oxide (30). Although it was suggested that OmcZ filaments might account for the high conductivity of anode biofilms (13), this hypothesis is inconsistent with the poor current production by strain Aro-5 and the low conductivity of its biofilms (18). Furthermore, OmcZ is localized near the anode-biofilm interface, OmcZ filaments are not observed coursing through the bulk of the biofilm (33).

In conclusion, eliminating artifacts by directly examining filaments emanating from cells has demonstrated that *G. sulfurreducens* expresses e-pili in abundance, consistent with multiple lines of evidence from previous studies (10, 11) that have indicated that *G. sulfurreducens* e-pili are an important component in long-range extracellular electron transport. The cells examined produced few OmcS-based filaments. The physiological significance of cytochrome-based filaments is yet to be determined.

## Methods

### Culture source and growth conditions

The strain of *G. sulfurreducens* expressing wild-type PilA pilin gene as well as strain Aro-5, which expresses a synthetic pilin gene designed to yield poorly conductive pili, and strain ΔomcBESTZ, which features deletions in five major outer-surface *c*-type cytochromes, were obtained from our laboratory culture collection and were previously described in detail (18, 24). Cells were grown in medium with acetate (10 mM) as electron donor and fumarate (40 mM) as electron acceptor as previously described (34). Cultures for AFM analysis were grown in the acetate-fumarate medium at 25 °C, a condition known to promote expression of e-pili (16).

### Analysis with atomic force microscopy

An aliquot (50 μl) of culture was drop cast onto a silicon wafer coated with a 35 nm layer of platinum, prepared as previously described (35). After 12 min, excess liquid was removed with a pipette and the substrate was washed twice with 50 μl of deionized water. Excess water was absorbed with filter paper and the preparation was allowed to air dry. Samples were equilibrated at 40% humidity inside scanning chamber of a Cypher ES, atomic force microscope (Asylum Research, Oxford Instrument) for at least 1 h at 25 oC. The filaments were first observed with tapping mode (AC-air topography) under repulsive force with a Pt/Ir-coated tip (PtSi-FM, NanoWorld AG) at a ~2.0 N/m spring force constant and ~70 kHz resonance frequency.

The conductance of individual filaments was determined in contact mode (force 30 nN) with the Pt/Ir-coated tip functioning as the translatable top electrode. Quadruplicate amplitude of ±0.4 V voltage at 0.99 Hz frequency was applied to get ca.8000 points per measurement. Three independent points from three individual wire (biological replicates) were analyzed to determine the conductance. Conductance was calculated from the linear slope between −0.2 to 0.2 V followed with the equation: Conductance = Current/Voltage as preciously described (23).

