## Supplemental material for "Direct Observation of Electrically Conductive Pili Emanating from *Geobacter sulfurreducens*"

**Supplemental Figure 1.** AFM height images corresponding to amplitude images of the primary text. (A) AFM height image of filaments emanating from *G. sulfurreducens* with the wild-type pilin gene (corresponding to Fig. 1A). (B) AFM height image of higher magnification of the region highlighted in the dashed frame in panel A (corresponding to Fig. 1B). (C) AFM height image of filaments emanating from *G. sulfurreducens* strain Aro-5 (corresponding to Fig. 2A). (D) AFM height image of strain Aro-5 at higher magnification illustrating the two filament types with yellow and white dashed lines designating cross sections for the 3 nm and 4 nm diameter filaments, respectively (corresponding to Fig. 2B). (E) AFM height image of filaments emanating from *G. sulfurreducens* strain △omcBESTZ (corresponding to Fig. 2E). (F) AFM height image at higher magnification showing 3 nm diameter filaments emanating from cell of strain △omcBESTZ (corresponding to Fig. 2F).

**Supplemental Figure 2.** AFM amplitude image of filaments emanating from *G. sulfurreducens* with the wild-type pilin gene in Sample 1, designating the six regions used to count the number of 3 nm and 4 nm diameter filaments. Height profiles were determined from the corresponding height images, which yield profiles similar to those shown in Supplemental Figure 1.

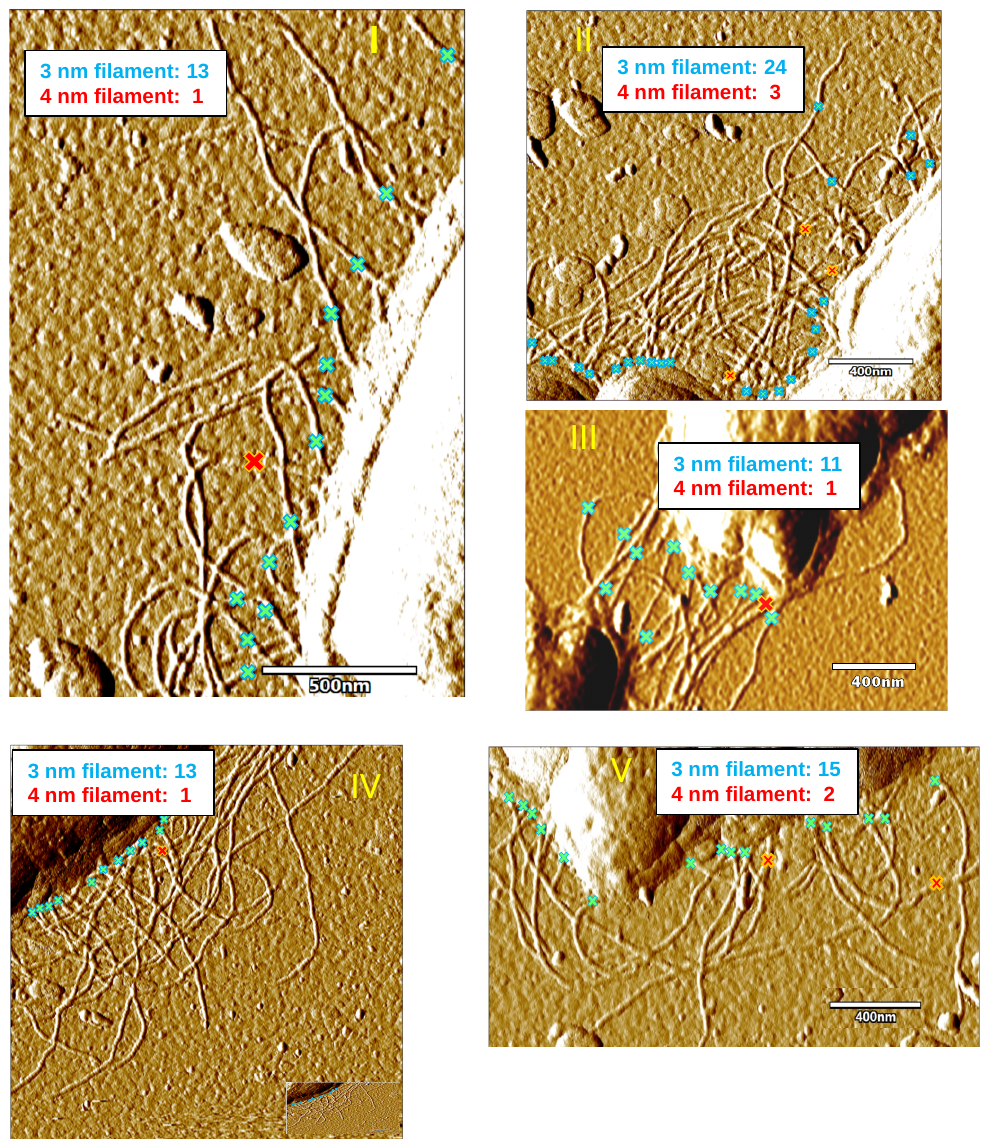

**Supplemental Figure 3.** Higher magnification of 5 regions noted in Fig. S2. Crosses designate the counted 3 nm (blue) and 4 nm (red) diameter filaments. Height profiles were determined from the corresponding height images, which yield profiles similar to those shown in Supplemental Figure 1. Insets show the results for each section.

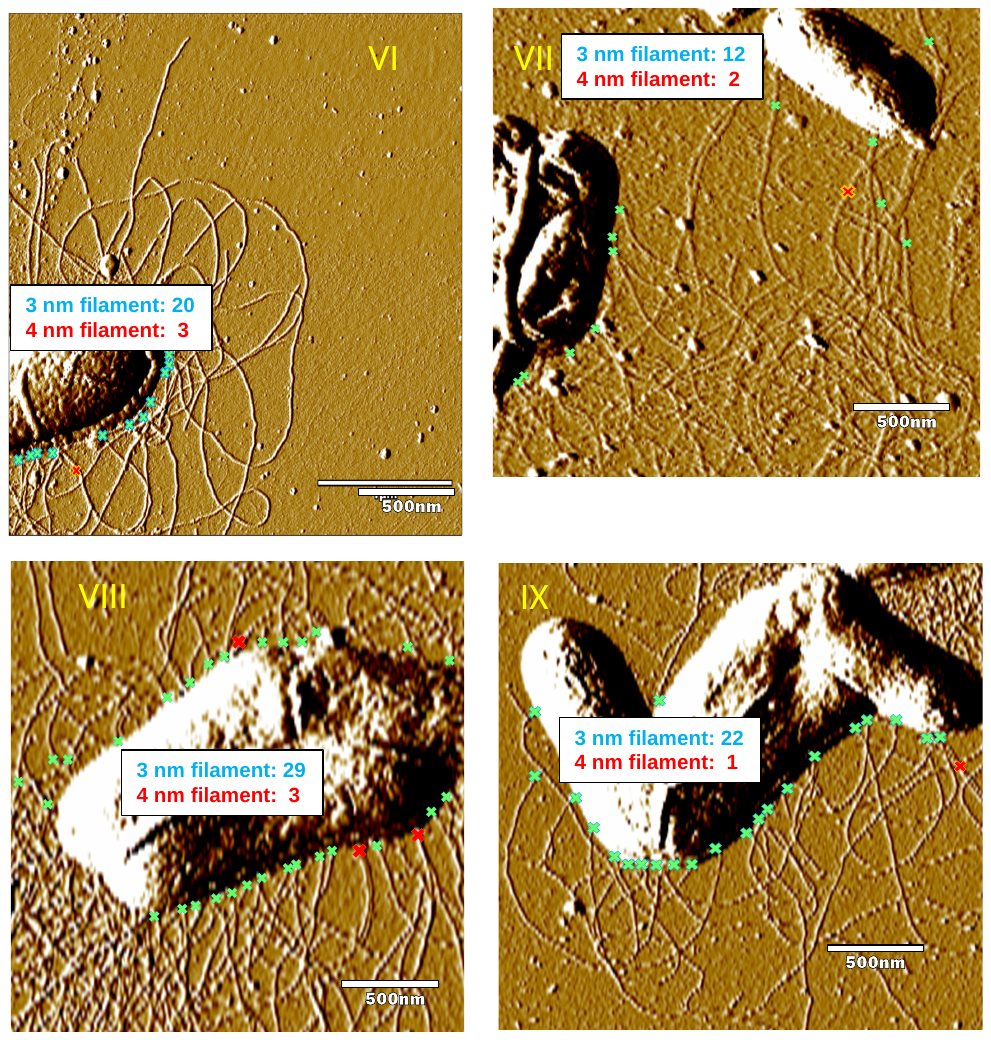

**Supplemental Figure 4.** AFM amplitude image of filaments emanating from *G. sulfurreducens* with the wild-type pilin gene in Sample 2 (designated regions VI and VII) and Sample 3 (designated regions VIII and IX). Crosses designate the counted 3 nm (blue) and 4 nm (red) diameter filaments. Height profiles were determined from the corresponding height images, which yield profiles similar to those shown in Supplemental Figure 1. Insets show the results for each section.

**Supplemental Figure 5.** Point-mode current response (I-V) spectroscopy measurements and conductance calculations of the three independent 3 nm diameter filaments from *G. sulfurreducens* with the wild-type pilin gene shown in Fig. 1D of the primary text. Three independent filaments are shown as row a, b, and c. Three independent measurements on each filament are shown as column 1, 2 and 3.

**Supplemental Figure 6.** Point-mode current response (I-V) spectroscopy measurements and conductance calculations of the three independent 4 nm diameter filaments from *G. sulfurreducens* with the wild-type pilin gene shown in Fig. 1D of the primary text. Three independent filaments are shown as row a, b, and c. Three independent measurements on each filament are shown as column 1, 2 and 3.

**Supplemental Figure 7.** AFM amplitude image of filaments emanating from *G. sulfurreducens* strain Aro-5 in Sample 1 (Regions I and II) and Sample 2 (Regions III and IV). Crosses designate the counted 3 nm (blue) and 4 nm (red) diameter filaments. Height profiles were determined from the corresponding height images, which yield profiles similar to those shown in Supplemental Figure 1. Insets show the results for each section.

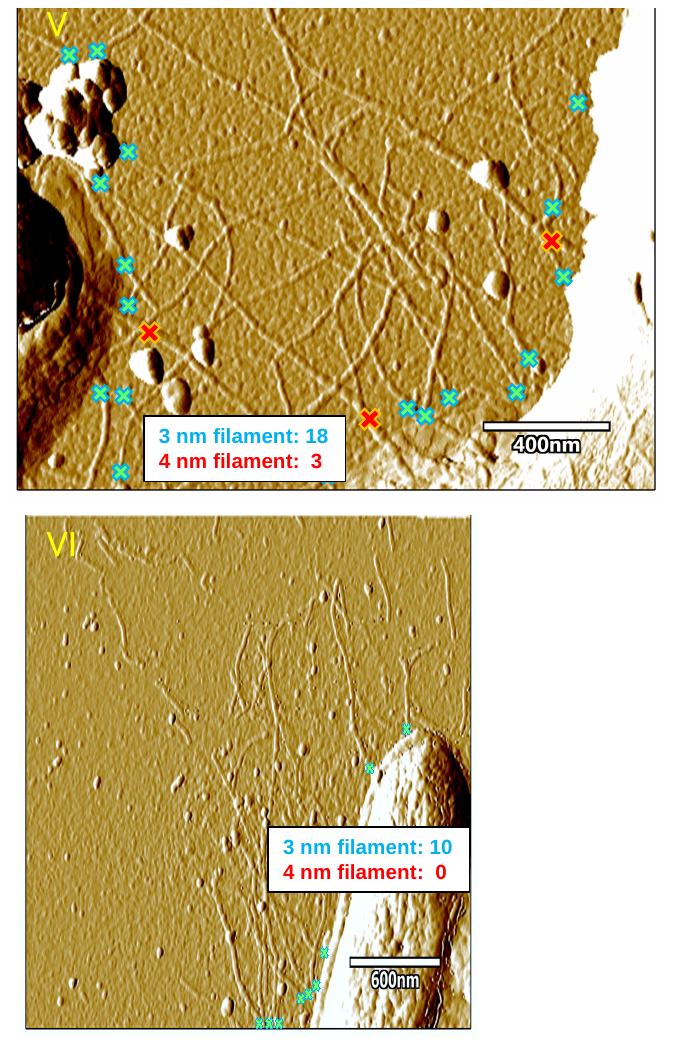

**Supplemental Figure 8.** AFM amplitude image of filaments emanating from *G. sulfurreducens* strain Aro-5 in Sample 3, Regions V and VI. Crosses designate the counted 3 nm (blue) and 4 nm (red) diameter filaments. Height profiles were determined from the corresponding height images, which yield profiles similar to those shown in Supplemental Figure 1. Insets show the results for each section.

**Supplemental Figure 9.** Point-mode current response (I-V) spectroscopy measurements and conductance calculations of the three independent 4 nm diameter filaments from *G. sulfurreducens* strain Aro-5 shown in Fig. 2D of the primary text. Three independent filaments are shown as row a, b, and c. Three independent measurements on each filament are shown as column 1, 2 and 3.

**Supplemental Figure 10.** Point-mode current response (I-V) spectroscopy measurements and conductance calculations of the three independent 3 nm diameter filaments from *G. sulfurreducens* strain Aro-5 shown in Fig. 2D of the primary text. Three independent filaments are shown as row a, b, and c. Three independent measurements on each filament are shown as column 1, 2 and 3.

**Supplemental Figure 11.** Point-mode current response (I-V) spectroscopy measurements and conductance calculations of the three independent 3 nm diameter filaments from *G. sulfurreducens* strain △omcBESTZ shown in Fig. 2F of the primary text. Three independent filaments are shown as row a, b, and c. Three independent measurements on each filament are shown as column 1, 2 and 3.
